# Computational Tools for the Analysis of Meiotic Prophase I Images

**DOI:** 10.1101/2023.11.15.567191

**Authors:** James H Crichton, Ian R Adams

**Affiliations:** University of Exeter; MRC Human Genetics Unit

**Keywords:** Meiosis, bioimage analysis, segmentation, axis, recombination, chromosome

## Abstract

Prophase I is a remarkable stage of meiotic division during which homologous chromosomes pair together and exchange DNA by meiotic recombination. Fluorescence microscopy of meiotic chromosome spreads is a central tool in the study of this process, with chromosome axis proteins being visualised as extended filaments upon which recombination proteins localise in focal patterns.

Chromosome pairing and recombination are dynamic processes, and hundreds of recombination foci can be present in some meiotic nuclei. As meiotic nuclei can exhibit significant variations in staining patterns within and between nuclei, particularly in mutants, manual analysis of images presents challenges for consistency, documentation, and reproducibility. Here we share a combination of complementary computational tools which can be used to partially automate the quantitative analysis of meiotic images. 1) The segmentation of axial and focal staining patterns, to automatically measure chromosome axis length and count axis-associated (and non-axis associated) recombination foci; 2) Quantification of focus position along chromosome axes to investigate spatial regulation; 3) Simulation of random distributions of foci within the nucleus or along the chromosome axes to statistically investigate observed foci-axis associations and foci-foci associations; 4) Quantification of chromosome axis proximity to investigate relationships with chromosome synapsis/asynapsis; 5) Quantification of and orientation of focus-axis distances. Together these tools provide a framework to perform routine documentation and analysis of meiotic images, as well as opening up routes to build on this initial output and perform more detailed analyses.

## 1. Introduction

During meiotic prophase I homologous chromosomes are organised into chromatin loops, anchored into an axial pattern by cohesion complexes [1]. Further proteins assemble along the chromosome axis, including HORMAD proteins [2] and the axial element proteins of the synaptonemal complex [3–5]. Meiotic recombination is initiated by double strand DNA break formation early in meiotic prophase I, with the chromosome axis playing a central role in its regulation, as both a hub around which recombination events physically organise [6], and as a regulator influencing the distribution of recombination events [5]. Repair of these DNA double strand breaks drives pairing and synapsis between homologous chromosomes, which in turn is accompanied by completion of synaptonemal complex assembly. Fully assembled synaptonemal complex then holds homologous chromosomes in close proximity along the length of their axes while meiotic recombination progresses and exchanges genetic material between homologous chromosomes [5].

Fluorescence microscopy is a particularly valuable tool in the study of meiosis. The progressive formation of chromosome axes and pairing of homologous axes is used to define substages of meiotic prophase I progression, and the quantification of axis length can be used to indicate variation in chromatin loop/axis organisation [7]. Recombination foci, which are typically associated with chromosome axes, are often abundant, exceeding 200 per nucleus [5], and are commonly counted to monitor progression of recombination. Such scoring is typically performed manually, and can be subject to subjective scoring biases. As recombination intermediates can follow different repair pathways and result in different repair outcomes, variation between and within nuclei can be difficult to capture. More complex analyses such as quantifying the position of recombination intermediates relative to each other along chromosome axes, typically to measure crossover interference, are almost exclusively restricted to low abundance recombination intermediates partly due to it being impractical to measure the positioning of more abundant foci [8, 9]. More automated alternatives have however recently been developed which begin to accelerate this process [10].

Advances in microscope resolution have dramatically improved the ability to visualise intricate events taking place during meiotic prophase I, particularly enabling the resolution of the components of the synaptonemal complex, and the relative positioning of axis-associated proteins [11–14]. These gains in resolution present many new opportunities for quantitative analysis of images of meiotic cells, but the extraction of this data requires the development of computational tools which can be used reproducibly. Several such tools have been developed, particularly for the analysis of single molecule localization microscopy images (SMLM) [11–13], however there remains a lack of generally applicable tools to address many research questions.

Here we share a toolkit of open-source computational programs which can be used in the analysis of meiotic prophase I chromosome spread images. These tools provide an efficient and effective framework for recording image data, improving the speed, documentation, and reproducibility of analysis, and generate a system of metadata which can be further explored, enabling greater insight into the complex processes taking place along chromosome axes during meiotic prophase I.

## 2. Materials

The analytical tools described, are designed to run in Fiji (ImageJ) 1.53f [15] and Python 3.10.9, both of which can be freely downloaded (https://fiji.sc/, https://www.python.org/downloads/). Code was written on a Windows operating system. Fiji scripts require the installation of the “IJPB-plugins” [16], “Neuroanatomy” [17], and “SCF MPI CBG” plugins, which can be added by selection within the Help>>Update>>Manage Update Sites setting in Fiji. All code described for the analysis of meiotic images can be downloaded from https://github.com/JamesCrichton/Meiotic-Image-Analysis-Toolkit. as a zip file, and extracted to the Desktop (Note 1). All code herein is designed to process stacked 2D tiff images of chromosome spreads. These preparations are beneficial for analysing chromosomes and associated proteins as nuclei are typically spread out and quite flat. Where z-stacks are taken to capture signal in 3D, we reduce images to 2D by maximum intensity projection before analysing. Example images and metadata are included in the zip file. To run the Image Segmentation macro first open Fiji then simply drag and drop this file into the Fiji toolbar. A stack image to analyse should be open before running the script, this can similarly be dragged and dropped into the toolbar (test image provided: “Example_Pachytene_DAPI_RPA_SYCP3.tif”).

Positioning of focal staining on chromosome axes, quantification of paired axis proximity and positioning of foci relative to axial pairs requires tracing of axes using SNT (in Neuroanatomy plugin). Guidance on using the SNT plugin can be found here: https://imagej.net/plugins/snt/step-by-step-instructions. The subsequent interpretation of saved .traces files in our scripts uses the TracePy package by Ziwei Huang. This should be downloaded from https://github.com/huangziwei/TracePy as a zip file and similarly extracted to the Desktop (Note 1). Example scripts using python functions to batch process files are written with the assumption that these repositories are located on the user’s Desktop. Python scripts will require modification to direct towards users’ specific images. We edit and run these scripts using the Spyder IDE (integrated development environment) downloaded with Anaconda (https://www.anaconda.com/). Measurements generated from all images are not scaled, but given in pixel values. Images analysed in the data presented are previously studied synapsed pachytenes (*Tex19.1*^+/+^, *Tex19.1*^+/-^, *Tex19.1*^-/-^) [18] images by widefield microscopy, and structured illumination microscopy images of *Sycp1*^+/-^ and *Sycp1*^-/-^ spermatocytes from other publications [19, 20] (slides kindly provided by Dr Willy Baarends [21]).

## 3. Methods

### 3.1 Axis length measurement and axial focus counting

Cell staging, counting of axial foci, and measurement of axis length are common analyses conducted on meiotic prophase I images. Here we share an Fiji macro which guides users to record the prophase I substage of their image and any comments. The macro then segments and measures the chromosome axes, segments and counts foci, and finally measures the proximity of foci to axes is measured.

Begin by opening an image to be analysed (Note 1. Note that when using OneDrive, a OneDrive Desktop is the system default. This location is not the default for our program. Rather, it is the user Desktop.

Note 2). This should be 2D and contain a minimum of two channels (axis and foci staining). The example image in the “Widefield_sample_data” folder, “Example_Pachytene_DAPI_RPA_SYCP3.tif” is a mouse pachytene spermatocyte with stained axial elements (SYCP3), recombination foci (RPA), and nucleus (DAPI). Nuclear staining is not essential for the script to run but provides a useful guide to select the region of an image to analyse. This image was captured on a diffraction-limited widefield microscope, so the separate paired axial elements cannot be resolved.

Once you have your image open, run the “Image Segmentation.ijm” macro in Fiji (Figure 1, Video 1). The user will be guided through a dialogue to define the channel of the stack image corresponding to the axial and focal staining to be analysed. Distinct axis and foci staining patterns are expected to be in different channels in the image to be analysed. If needed, the user can open the Fiji Channels tool at this point to inspect the different channels in the image. If an image has previously been analysed then segmented staining patterns can be reapplied, enabling the addition and comparison of further focal or axial channels (Note 3). To facilitate this type of analysis, the user can select the ‘Pre-existing mask’ and ‘Pre-existing labelmap’ options for the axis and foci channels respectively. The user also has an option to customise the titles of their analyses, which will be used in the output filenames, however unless the user is re-analysing or performing additional analysis on a previously analysed image we recommend keeping these titles as their defaults.

**Figure 1.**
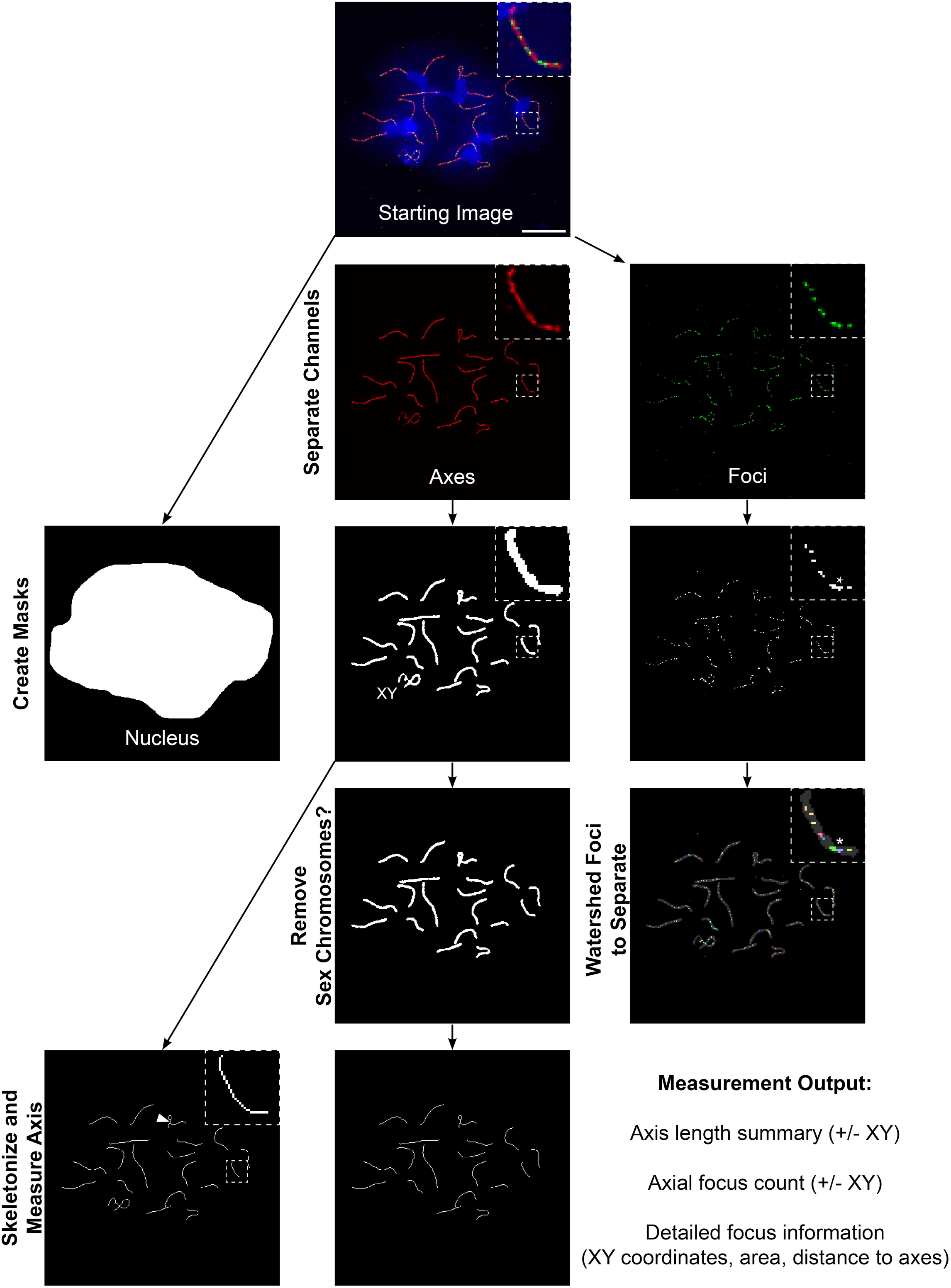
Segmentation pipeline. Beginning with a fluorescence image, nuclear, axial, and focal binary masks are generated. Sex chromosomes (marked “XY” on Axes Mask image) can be erased to create an autosomal mask. Axial masks are skeletonized to create single-pixel thick lines, the length of which are automatically measured. Occasionally, proximal axes can become fused during this process (arrowhead), underestimating length measurements. Adjacent points fused together in the focus mask (asterisk) can be separated using a maxima watershed function. 10μm scale bar.

The user will be asked to define the destination of the analysis output. This output will consist of a folder of metadata named in the format “*imagename*_Output”. We recommend arranging the images to be analysed for an experiment or condition into one folder before starting the analysis, then selecting this folder as the output destination for these new metadata folders generated. This organisation of images and metadata folders is assumed in subsequent analytical pipelines. If the image has been previously analysed and the data has been saved in this format, segmented regions of interest (ROIs) and an “Image_Summary.csv” file will be opened automatically. If the image has not been analysed before, a dialog will open asking the user to define the cell’s meiocyte sub-stage, add any comments, and define whether the sex chromosomes should be separated from analysis, using the “Separate Autosome Scoring” checkbox. In mammals, recombination and synapsis behave differently on heterologous XY sex chromosomes than they do on the autosomes [22], therefore it can be important to analyse recombination and synapsis on the autosomes independently from the sex chromosomes (Note 4). If this option is chosen then analysis is run on all axes, as well as just the autosomal axes (with the identified sex chromosomes removed).

Image segmentation begins with the nucleus, the user should use the ‘Selection tools’ in the Fiji toolbar to draw round the nucleus/area to be analysed. A freehand tool is selected by default. The nucleus is typically visualised in meiotic samples by staining with DAPI or other DNA-binding fluorescent stains. Defining the nuclear area is useful if an image contains multiple nuclei, and is also used when generating random nuclear distributions of foci for that image (section 3.5). This ROI will be labelled “Nucleus_Mask” in the ROI manager. Only signals found within this ROI will be analysed by the macro from this point on.

Chromosome axes are segmented next. In meiotic chromosome spreads, axes are typically labelled with synaptonemal complex components such as SYCP3 in mammals, though chromosome core components such as SMC3 cohesin could also be used (Note 5, Note 6). The macro first asks the user to adjust brightness/contrast to visualise the axes (these brightness/contrast adjustments purely to help the user visualise their image. Changes should not be “Applied” to avoid propagation into the analysis steps). The user can then experiment with applying a Gaussian filter to the axis channel to reduce noise and improve the quality of the segmentation (Note 7). The Gaussian filter will accept positive integer values, and a 0 value will apply no filtering. Images with more off-axis background staining and discontinuities/fragmentation of axis staining may benefit from higher Gaussian filter values to reduce both fragmentation and off-axis staining being interpreted as chromosome axes. The macro records the settings used for Gaussian filtering in the “Image_Summary” output to allow the user to reproduce analyses from the original image. The user then generates a binary axial mask by setting a threshold above which the macro will consider staining to represent bone fide axis staining. This threshold is somewhat subjective, though there are auto-threshold methods available within the Fiji thresholding tool to reduce subjectivity. The resulting mask from the axes will be recorded in the ROI Manager and named with the given Axes analysis title (“Axes_Mask” by default).

If the user previously selected the “Separate Autosome scoring” option, the user will then have the option to remove the sex (XY) chromosomes from the axes mask using the Eraser Tool (in Drawing Tools). During this erasure process, the mask is overlayed on the original axial channel for reference and the two can be toggled on/off with the Channels Tool. The new axes mask will be have an “Autosome_” prefix in the ROI manager. The macro will count foci in the original mask, and in the autosome mask. If the user is choosing to reuse a previously generated mask, they will be invited to select this in the ROI Manager (Note 8).

The thickness of the chromosome axis staining will depend on the antibody used, the intensity of the immunostaining signal, and the image capture conditions. Therefore, reducing the axis mask to a single pixel thickness line along the middle of the axis can facilitate some types of analysis. To do this, the axis masks are skeletonized using an algorithm that removes pixels from the circumference of each axis in the mask until a continuous single pixel line remains. The quality of this process can vary depending on the axial mask used, therefore users are given the opportunity to manually refine the axis skeleton by removal or addition of necessary connections using the Eraser and/or Pencil drawing tools (set width to 1px) found in the Fiji toolbar (Note 9). Users can also comment on the quality of these skeletons in a subsequent dialogue for reference. The skeletonized axis mask will be labelled “Axes_Mask_Skeleton” in the ROI Manager.

A similar process of segmentation is followed for the foci channel. The macro will display an image with the axial mask outline overlayed on the foci for reference to aid foci segmentation. The macro first asks the user to adjust brightness/contrast to visualise the foci and identify areas with weak/closely spaced foci to monitor foci segmentation (these brightness/contrast adjustments are lost when the user proceeds, they are not propagated into the analysis steps). The user then generates a binary focal mask by setting a threshold above which the macro will consider staining to represent bone fide focal staining. This threshold is somewhat subjective, though there are pre-set auto-thresholding methods available within the Fiji thresholding tool to reduce subjectivity. In images with numerous foci it is often impossible to separate the foci by thresholding alone, so a maxima watershed step is included (Figure 1). The macro will ask users to set a maxima prominence which represents the minimum difference in intensity between surrounding pixels used to identify distinct peaks in regions with merged/overlapping signals. By default, the resulting focal masks will be labelled “Watershed_Foci_Total” in the ROI Manager, the positions of the foci maxima will be labelled “Watershed_Foci Maxima”, and a labelmap image of these foci will be saved as “Watershed_Foci.tif” which can be used in other analyses discussed later. These default names and related measurement titles will alter depending on the “Foci analysis title” set within the macro. The title of the axes mask will default to “Axes_Mask” and will similarly apply to the related files, ROIs and measurements, but can also be modified in the macro under “Axes analysis title”. Measurements relating to these individual foci are then automatically generated: area, circularity, centroid X/Y coordinates, distance from the nearest point on the axial mask, and distance from the nearest point on the autosomal mask (if selected). A basic summary of measurements is also recorded in the “Image_Summary.csv” file, including a count of foci on axes, off axes and on autosomes (if selected), and the Euclidean length of the skeleton masks produced (NB. All distance measurements produced are in pixel units). The mask ROIs in the ROI Manager are also saved as a RoiSet.zip file. If images are being reanalysed, the summary metadata is appended to the pre-existing “Image_Summary.csv” file each time whereas a new file is generated for the measurements of each focal and axial pattern analysed.

### 3.3 Validating counts from segmentation pipeline

To assess the reliability of the measurements generated using the “Image_Segmentation” macro we first investigated its ability to measure chromosome axis length. The autosomal axis length of a library of pachytene spermatocyte images was first measured using the SNT plugin [17]. This is a plugin which allows users to trace axial signals and is widely used for chromosome axis tracing in meiosis research [14, 23]. The image library consists of wild type, *Tex19.1*^+/-^, and *Tex19.1*^-/-^ mouse pachytene spermatocytes pooled together from a study into pachytene progression in this mutant [18]. The same images were also processed using the “Image_Segmentation” macro, generating automated axis length measurements. Fitting a linear model to the data for each image generated using these two methods gave an R^2^ of 0.93 demonstrating that the automated measurements capture a large amount of the variation in the manual measurements of axis length and performs to a very high standard. Drawbacks to the use of skeletonization to measure axis length include the systematic removal of pixels from object ends, and the potential fusion of adjacent regions of axes during thresholding resulting in multiple axes being reduced to single skeleton traces. Both these caveats reduce the total axis length measured (Figure 1, Figure 2A). This reduction likely accounts for the slight underestimate of length using the automated method (coefficient = 0.87) and should be considered during the segmentation process.

**Figure 2.**
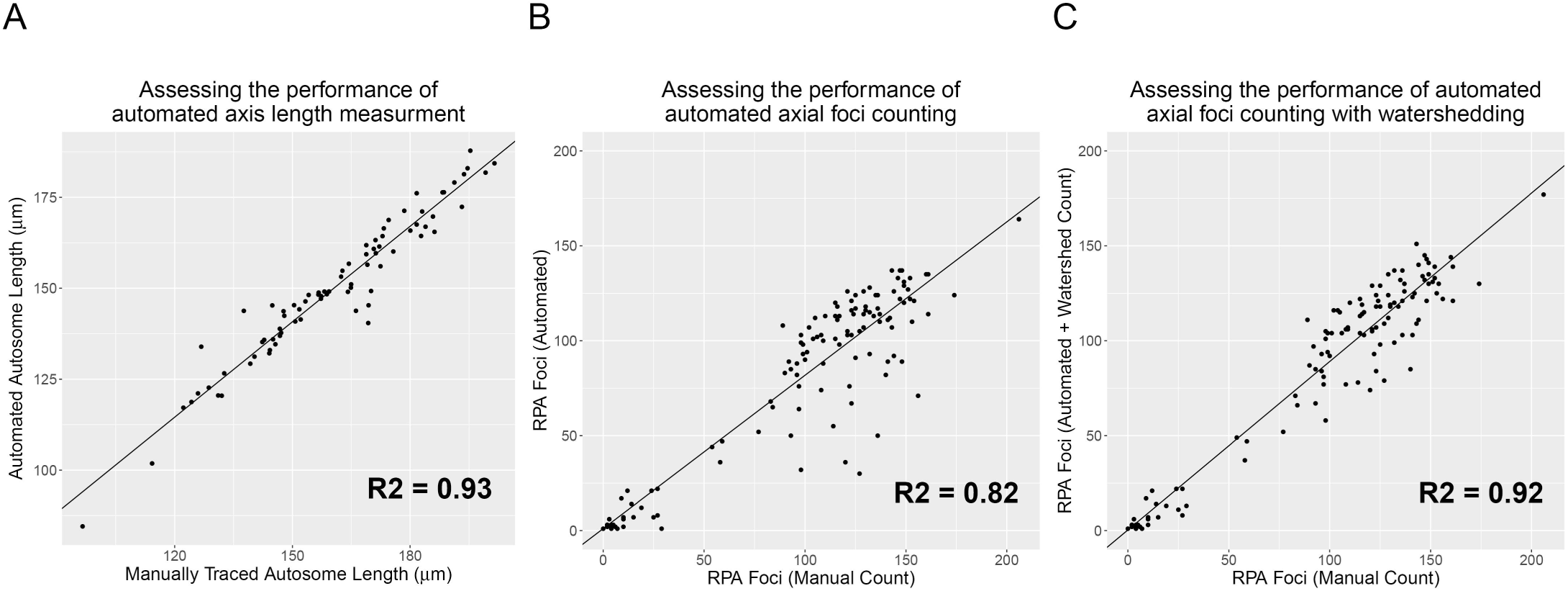
Automated measurement validation. A) Scatter plot comparing total autosomal lengths for a library of images measured by manual tracing using SNT, and automated measurement using the Meiosis-Toolkit Image_Segmentation macro (n=78, R^2^ =0.93, coefficient=0.87, intercept=9.57). Pixel output of analysis is converted to microns using original image-capture metadata. B) Scatter plot comparing total autosomal RPA foci in a library of images measured by manual counting [18] and automatically, using the Meiosis-Toolkit Image Segmentation macro, without using watershedding to separate foci (n=120, R^2^ =0.82, coefficient=0.81, intercept=1.03), and C) with watershedding to separate foci (n=120, R^2^ =0.92, coefficient=0.89, intercept=0.33).

Similarly, the performance of the automated focus counting feature of the macro was assessed by counting autosomal RPA foci in a library of pachytene spermatocytes manually [18], and automatically using the macro. Initially the segmented focus masks were analysed directly, without watershedding. Comparison with the manual counts gave an R^2^ of 0.82 (Figure 2B), showing that a large amount of the variation identified by manual counting was recapitulated by the automated count. However, it was apparent that counts for many images were underestimated. The RPA foci counted are highly abundant, typically 100-150 per nucleus, and as such, separation by thresholding alone is often not sufficient, resulting in foci fusing together in masks. Applying the watershedding step greatly improves performance of the macro, producing an R^2^ of 0.92 (Figure 2C), demonstrating a high performance even when counting abundant foci.

### 3.4 Focus position on axis

Analysis of focus position along the chromosome axis is a measurement often generated for the crossover events which form well established spatial pattens along the 1-dimensional axis. Similar patterns have also been reported for more abundant early recombination markers [9] but such measurements have rarely been performed [23], and analyses have been restricted to a selection of chromosomes. This positional measurement can be a laborious and manually involved process, poorly suited to these more abundant axial foci. Here we present a method to rapidly automate the calculation of this positional data (Figure 3).

**Figure 3.**
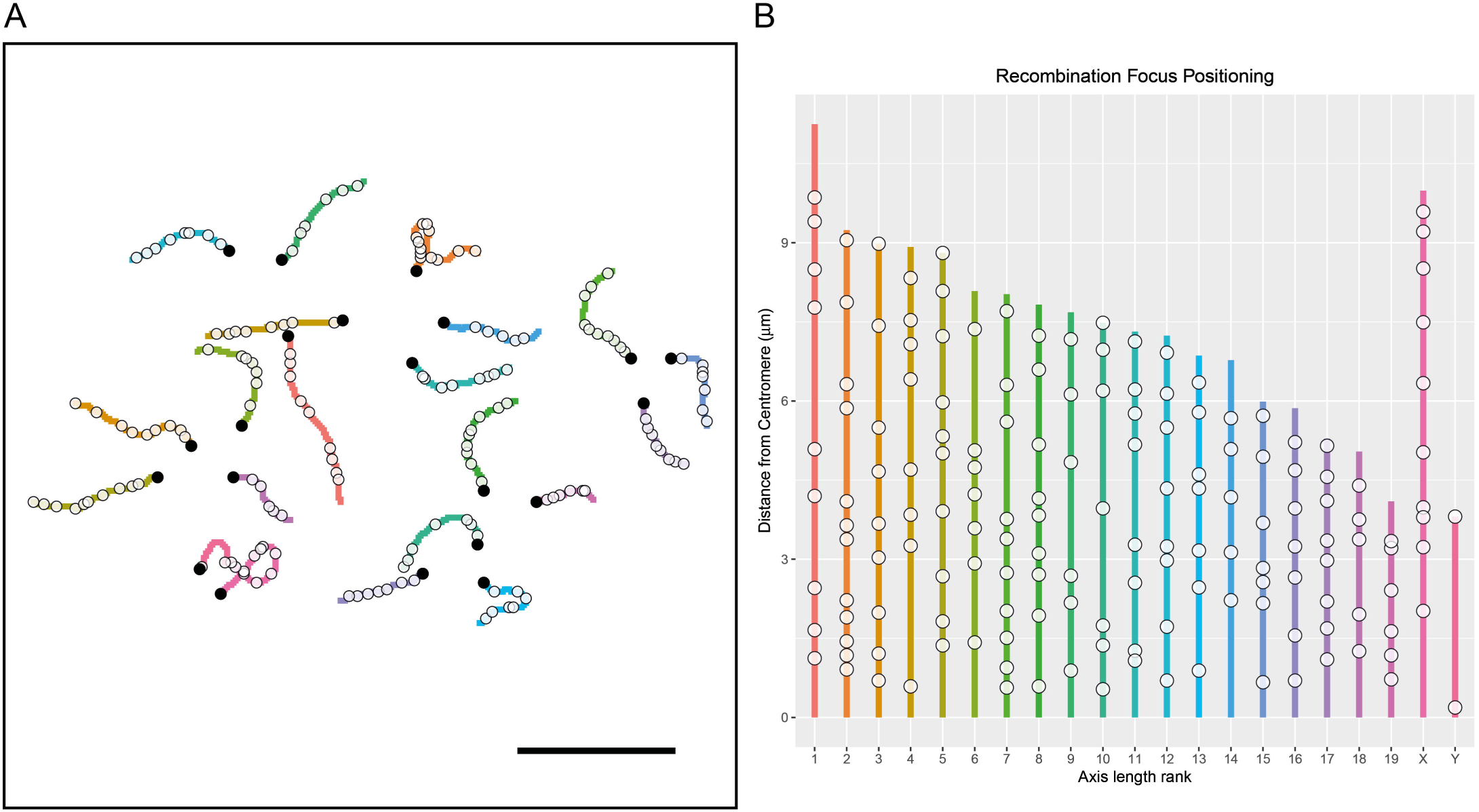
Focus positioning on chromosome axes. A) Plot displaying the 2D organisation of chromosome traces as in the original image, orientated with centromeres marked in black circles. Opaque circles mark the centroids of segmented RPA foci selected for overlap with the segmented axis mask. 10μm scale bar. B) 1D plotting of chromosome traces and associated positioning of the segmented foci. Chromosome number estimated by axis length ranking. Distances converted from pixel values to μm using image capture metadata.

First the image should be segmented using the Image_Segmentation macro, this will produce a labelmap image for the segmented foci (default name “Watershed_Foci.tif”). To calculate the position of each segmented focus along the chromosome axis, the axes must also be traced using SNT, an Fiji tool which forms part of the Neuroanatomy plugin [17]. A guide to the installation of SNT and its use is included in the Materials section. The raw image being analysed should be used as the template to generate these traces, with the axes and DAPI channels visible and the “Tracing channel” in SNT set to the channel with axial signal (Note 10, Note 11). Studying mice, which possess acrocentic chromosomes, we begin each trace at the DAPI-bright centromeric end of the axis to orientate the data. An alternative approach that would be applicable to non-acrocentric chromosomes would be to include a centromeric marker, which can be segmented as an axial focus in the Image Segmentation macro, and its position along traced axes subsequently calculated. This position can be used as a reference point for users to manually recalculate measurements to. A further alternative is to trace from each centromere and rename the traces differently in SNT for each arm (e.g., 1p and 1q), these names will be preserved in the processing. Similarly, naming of sex chromosomes in these data can also be beneficial. Save the traces in the image’s metadata folder as a “.traces” format to ensure it is identified in subsequent scripts in this toolkit.

An example batch analysis can be found in the “Focus_positioning_on_traces.py” file within “Example_Python_Scripts”. Open this file in a text editor or using your preferred integrated development environment (IDE), and modify the paths to locate the necessary modules and images. The defaults are set to run example data and assume everything has been extracted to the desktop (Note 1). The position of each focus along its closest traced axis will be added to the existing data relating to each focus, producing a file ending “_and_positioning.csv” in the metadata folder.

### 3.5 Comparing observed focus positioning to random distributions

To explore whether observed localisation patterns of segmented foci relative to axial or other focal segmented staining patterns reflect genuine associations, it can be useful to compare these measurements to those generated if the foci were randomly distributed [11, 19, 20]. The “Nuclear_focus_shuffling.py” script, in the “Example_Python_Scripts” folder, selects the foci in an image, filters out foci that are very small or very large, then assigns each focus in that image to a random location within the nucleus (defined by the nucleus mask) and determines the relative position of each shuffled focus and the axes. The example file for this script stored in “Widefield-sample_data”. The shape of each focus remains intact during this process, its centroid is repositioned, and new focus positions do not overlap other foci or extend beyond the nuclear boundary (Figure 4). The script performs multiple rounds of shuffling to generate a set of shuffled distributions of foci that can be compared with the observed dataset. Before running the script, paths to modules included in this package, paths to the image metadata and the file names of segmented focal and axial masks to be analysed, thresholds for foci filtering and the number of shuffles need to be set according to the instructions in the script itself. The resulting random distributions can be used to determine whether foci are more frequently associated with axis, or closer to the axes, in each image than might be expected by chance [19, 20]. In addition to generating these measurements, the example script also saves images of the size-filtered segmented foci selected for processing (“Size-selected_foci_labelmap.tif”), and one representative shuffled-foci image (“Shuffled_foci_labelmap.tif”).

**Figure 4.**
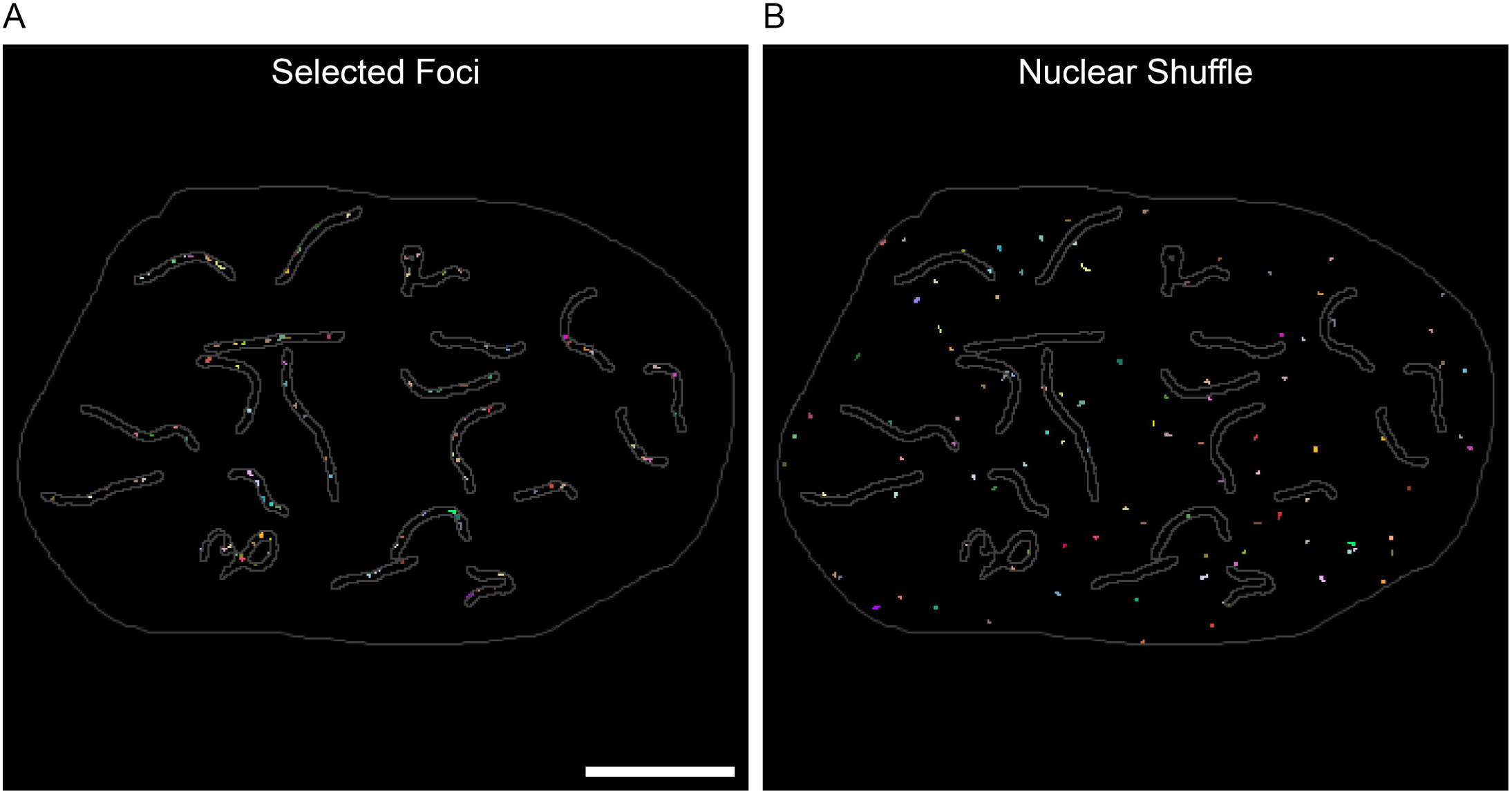
Nuclear focus shuffling. A) Segmented foci in their original positions. Separate foci coloured distinctly. Nuclear and axial mask outlines in grey. 10μm scale bar. B) An example image showing foci from (A) after shuffling throughout the nucleus.

When studying the proximity of two axis-associated proteins, it can also be useful to compare their observed proximity relative to that expected from each of these proteins being distributed along the chromosome axes at random [11]. A slightly different algorithm to the nuclear shuffling is required to achieve random shuffling along the chromosome axes as the edges of foci often extend beyond the edge of an axial mask. The “Axial_focus_shuffling.py” example script performs a similar process to that described for nuclear shuffling except that one set of foci centroids are randomly shuffled within that image’s axes mask rather than its nuclear mask. In addition, the foci are permitted to extend outside the boundary of the axial mask to better reproduce their natural distribution (Figure 5). To measure distances between two different foci labelled with different antibodies (A and B), distances from each shuffled B focus to the closest reference A focus are then measured (Figure 5F, Note 7). As well as generating these measurements, the example script also saves images of the size-filtered segmented foci selected for processing, and one representative axis shuffled-foci image.

**Figure 5.**
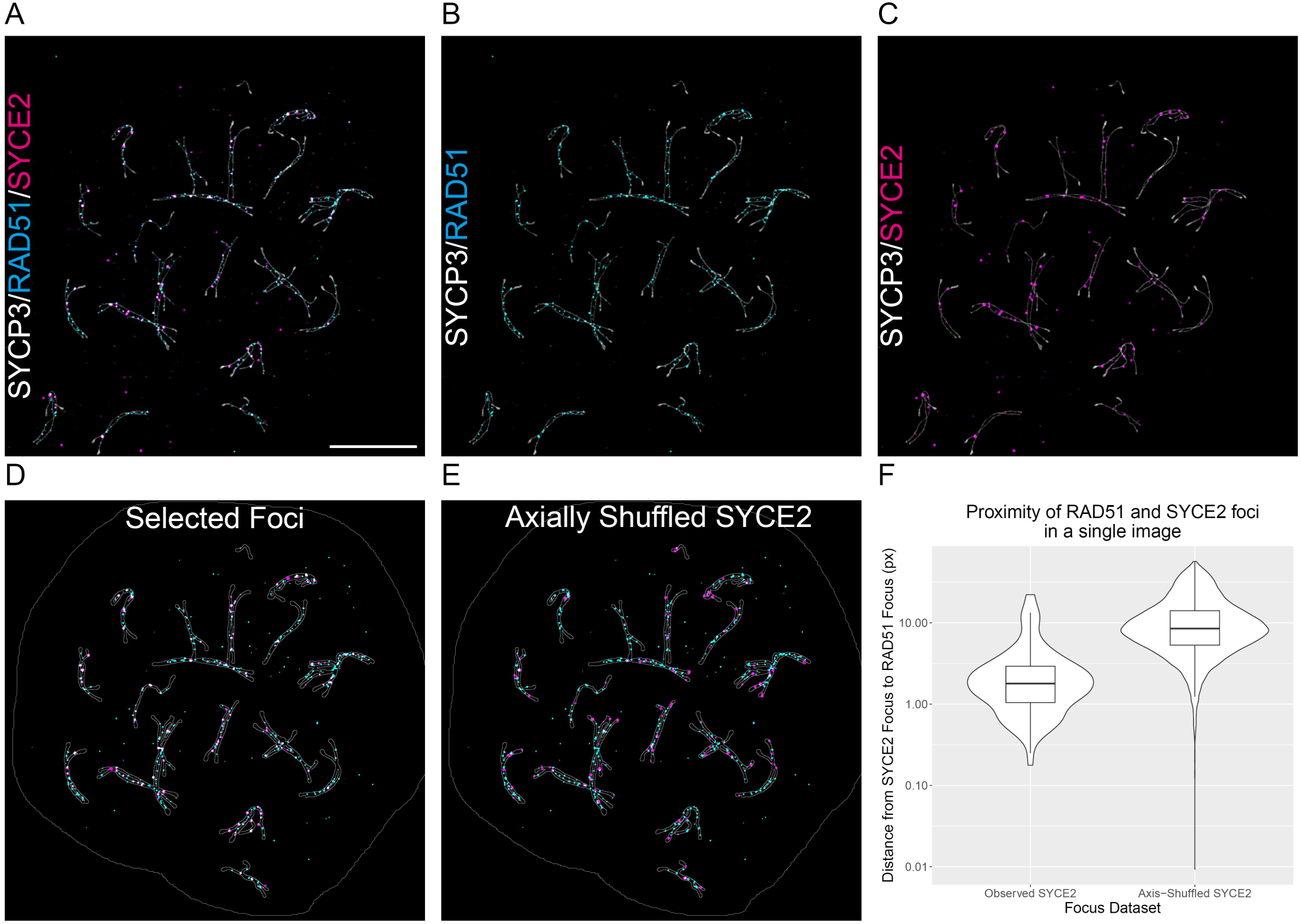
Axial focus shuffling. Structured illumination microscopy (SIM) images of *Sycp1*^-/-^ spermatocytes stained for the axial element protein SYCP3 (grey), and focal patterns seen by RAD51 (cyan) and SYCE2 (magenta) in this mutant. A) Merged original image, 10 μm scale bar, B) SYCP3 and RAD51 only, C) SYCP3 and SYCE2 only. D) Segmented RAD51 (cyan), and SYCE2 (magenta) foci in their original locations and E) shuffled around randomly the chromosome axis. Axial and nuclear masks outlined in grey. F) Violin plot demonstrating the focus distance measurements generated from this individual image.

### 3.6 Paired axis proximity

Super resolution imaging has made it apparent that homologous chromosome axis proximity varies along the length of the synaptonemal complex (Figure 6A, B). Quantification of these changes in proximity may reveal differences in synaptonemal complex structure and/or chromosome interactions. Pairing of chromosome axes is also seen in some synapsis-defective cells such as synaptonemal complex mutants, where quantification of pairing could help to better understand and stratify different asynapsis phenotypes [19, 20].

**Figure 6.**
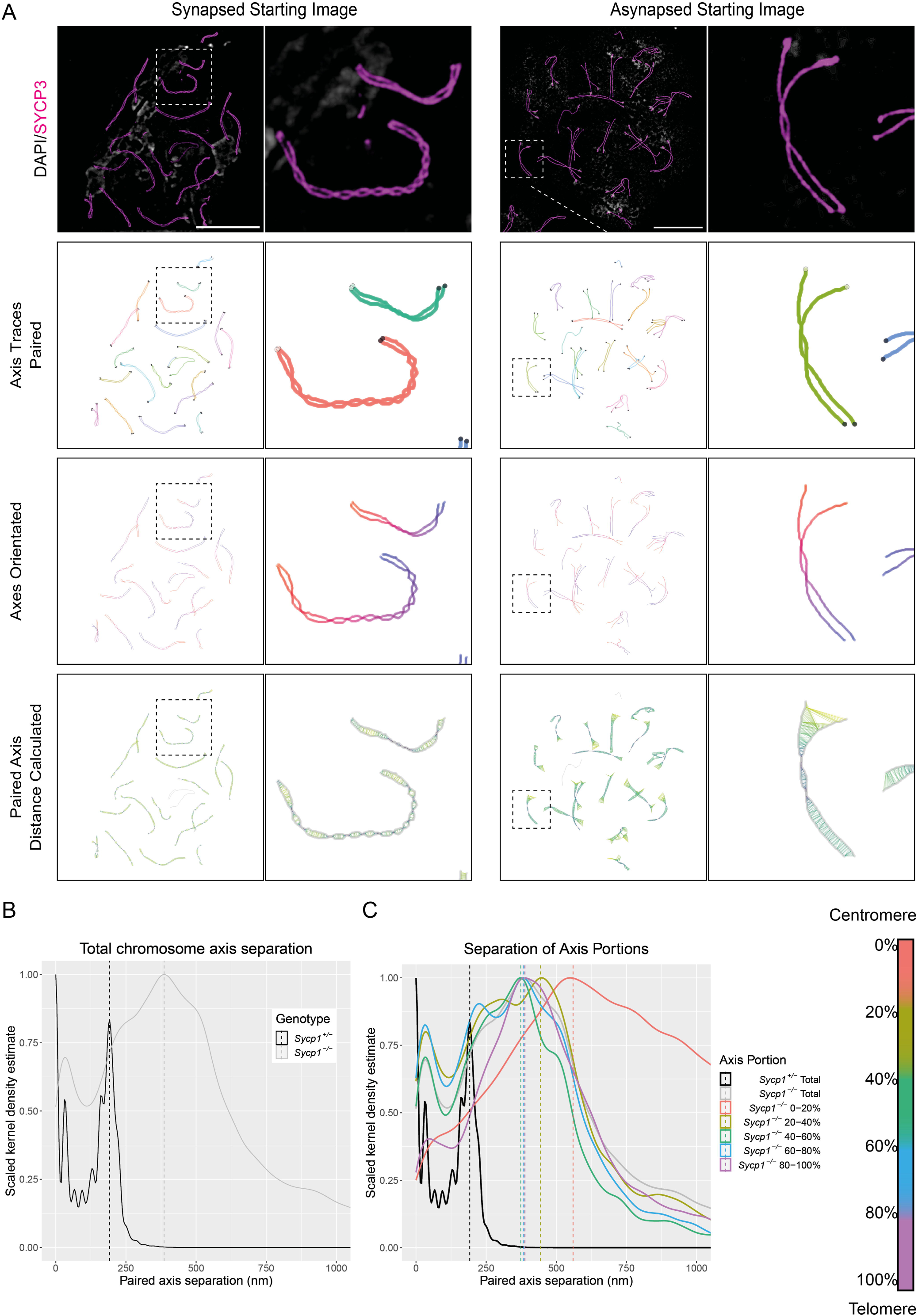
Homologous axis proximity. A) Synapsed (*Sycp1*^+/-^) and asynapsed (*Sycp1*^-/ -^) pairs of homologous chromosomes in spermatocytes stained with DAPI (grey) and antibodies to SYCP3 (magenta) and imaged using SIM. 10μm scale bar. Pairs of traced axes are recognised (colours paired) and orientated (centromere – red, to telomere – blue), and the minimum distance between each axis trace pixel and its partner axis is calculated (connections drawn and coloured dark blue (close) to yellow (distant)). Distance colouring scales are normalised separately to each image. B) Kernel density plot for paired axis separation for total traced axes from synapsed *Sycp1*^+/-^ spermatocytes (9 images of synapsed pachytene spermatocytes from 2 mice, including 171 pairs of traced axes) and asynapsed *Sycp1*^-/-^ spermatocytes (2 mice, 18 cells with a minimum of 18 pairs of traced axes each). Distribution of these distances (above zero) peak at 191nm (*Scyp1*^+/-^) and 386nm *Scyp1*^-/-^) respectively, indicated by dasher vertical lines. C) Kernel density distribution of paired axes separation in five equal portions of each *Sycp1*^-/-^ axis length. Dashed vertical lines indicate the peak separation for each group (NB. Distances have been scaled from the original pixel data into nm using metadata from image capture). Peak separation of total *Sycp1*^+/-^ axes = 191nm, total *Sycp1*^-/-^ axes = 386nm, *Sycp1*^-/-^ 0-20% = 560nm, *Sycp1*^-/-^ 20-40% = 444nm, *Sycp1*^-/-^ 40-60% = 373nm, *Sycp1*^-/-^ 60-80% = 384nm, *Sycp1*^-/-^ 80-100% = 388nm. Schematic on the right indicates the separation of chromosome axes into five colour-coded sections.

We provide a tool to calculate axis proximity using chromosome trace input generated by the SNT plugin. Users must first use SNT to trace the individual chromosome axes from their original fluorescence image (Note 13), renaming each member of a pair of axes with a shared number and a different letter e.g. “1a” and “1b”. These traces should be saved as a .traces file in the image metadata folder. The script recognises the names of the chromosome pairs, and these are oriented assuming the start position of the traces is the DAPI-bright centromere. For each pixel in each axis, the closest corresponding pixel in its partner axis will be identified and the distance to it calculated (Figure 6A) (Note 14). This nearest neighbour calculation involves organising the coordinates of one axis into a k-dimensional tree to allow efficient querying, then using this to identify the pixel closest to each pixel in the partner axis. This process is repeated for each member of each axis pair.

An example script (“Paired_axis_proximity.py”) for performing this task in a batch of images can be found in “Example_Python_Scripts”. This script will process the example image data in “SIM_sample_data”. To use this script, open the file and modify the paths to locate the necessary modules and images (as before, the image path should be a folder containing the metadata subfolders for each image, containing a .traces file of axis traces). The defaults are set to run example data and assume everything has been extracted to the desktop. The script will generate a spreadsheet of data titled “Axis_Trace_Measurements.csv”, which will save into each metadata folder.

We share an example analysis in Figure 6 investigating the proximity of paired chromosome axes in image datasets of synapsed control pachytene spermatocytes and mouse mutants unable to form a synaptonemal complex due to deletion of the gene encoding the transverse filament protein SYCP1, but capable of distinctive chromosome pairing in a pachytene-like stage [21]. Calculating paired axis proximity in the synapsed *Sycp1*^+/-^ controls reveals the axis separation varies, with density plots for axis separation peaking at 191 nm likely reflecting separation of the traced centres of SYCP3 axes at sites of complete synaptonemal complex assembly, and also peaking at 0 nm, indicating that the synaptonemal complex at many sites measured is twisted or collapsed (Figure 6B). Separation of *Sycp1*^-/-^ axes is higher than controls, with density plots of SYCP3 axis separation peaking at 386 nm (Figure 6B), however it is clear from the images (Figure 6A) that axis proximity varies greatly across the length of the chromosomes. To investigate whether axis proximity differs is any particular region we split axes into 5 equal portions and asked how close axes are in each of these (Figure 6C). Axis proximity of most segments peaks in density plots within 60 nm of each other (384-444 nm), however far greater separation is seen in the most centromeric fifth of the axes, which shows a peak at 560 nm, 116 nm greater than the next highest section. Therefore, while pairing of chromosomes persists in the absence of synaptonemal complex formation in this mutant, the degree of pairing which is achieved is not uniform, with the centromeric ends of the chromosome displaying dramatically lower association with one another.

Homologous pairing of chromosomes in mice is promoted by recombination [24], therefore differences in pairing of chromosome regions could be related to regional differences in recombination frequency. *Sycp1*^-/-^ spermatocytes assessed for paired chromosome proximity were also co-stained for RAD51 (Figure 7A), and regional density of recombination foci can be investigated using the code described in section 3.6 (Focus Position on Axis). The position of RAD51 foci along all autosomes included in the analysis of axis proximity was calculated, pooled, and plotted, with axes segmented into 10% bins (Supplementary Figure 1). This reveals a consistent frequency of RAD51 foci along the centre of the axes (mean 11.3% of RAD51 foci in each of the 8 x 10% segments from 10-90%). However, strikingly fewer foci are positioned in the terminal 10% of the axes, with 7.9% of foci in the most telomeric 10% of the axis, and only 2.0% of RAD51 foci in the most centromeric 10% of the axes. This centromeric depletion of RAD51-marked recombination sites may be responsible for the lower axis proximity seen for the most centromeric 20% of paired axes (Figure 6C). This analysis also further highlights the value of the tools we are sharing and the data they generate.

**Figure 7.**
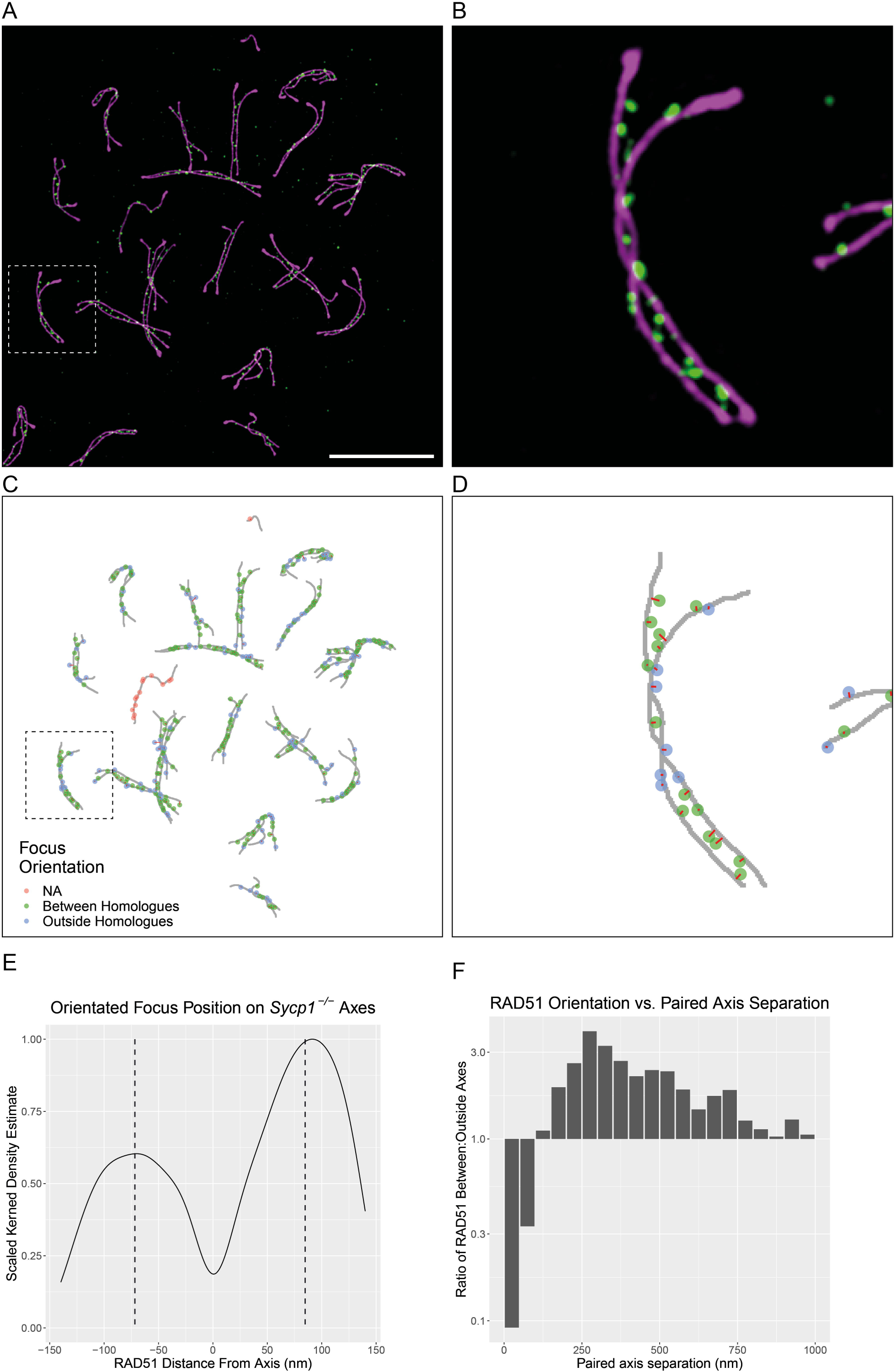
Focus positioning relative to individual homologous chromosome axes. A) Structured illumination microscopy (SIM) images of *Sycp1*^-/-^ spermatocytes stained for the axial element protein SYCP3 (magenta), and focal patterns seen by RAD51 staining (green). 10μm scale bar. B) Zoomed example of two paired homologous axes. C) Plotted coordinates of traced axes (grey) and the weighted centroids of segmented RAD51 foci coloured based on their orientation relative to their closest axis par (red=NA i.e. closest axis is not paired with its homologue; green = focus between homologous axes; blue = focus outside homologous axes). Red lines indicate the distance measured from the focus centroid to the closest pixel in the traced axis. D) Zoomed example axes. E) Kernel density estimate plot of RAD51 focus position relative to the centre of the closest axis trace. Focus distances orientated between paired axes are positive, distances away from a partner axis are negative. Only cells with a minimum of 18 pairs of traced axes are included (18 cells from 2 mice). Vertical dashed lines indicate the median positive and negative distances (Positive: 78.35μm, Negative: total = −72.92μm). F) Barplot showing the ratio of RAD51 focus orientation being between paired axes, to outside paired axes, against the separation of paired asynapsed chromosome axes in *Sycp1*^-/-^ spermatocytes, divided into 50nm bins.

### 3.7 Focus Positioning Relative to Homologous Axes

New levels of insight into the structural intricacy of events taking place during meiosis are being provided by super-resolution microscopy [11, 12, 14, 25]. It is important to be able to measure these spatial relationships to best utilise these advances in imaging technology. As well the existence of variation in proximity between homologous axes, the positioning of recombination foci within the paired chromosome axes also varies [25]. We share a method using a combination of axis trace data and segmented foci data to calculate the position of foci along the axes, their distance away from their closest axis, and whether the orientation of this position is between paired axes or outside of these.

To use this approach, focal staining patterns must first be segmented to generate a labelmap image. This can be done using the “Image Segmentation.ijm” macro described earlier. Paired axes must also be traced and named in the manner described for the calculation of paired axis proximity, with each member of a pair of axes having a shared name but different number e.g. “1a” and “1b”. These should be saved as a .traces file in the same image metadata folder which contains the segmented focus labelmap image among other data generated during segmentation. The proximity of axes must first be calculated using the “Paired_axis_proximity.py” script to generate an “Axis_Trace_Measurements.csv” file which will be read. The example python script, “Focus_position_relative_to_individually_traced_axis_pairs.py” will use this data and calculate the closest axis trace position for each segmented focus centroid, efficiently searching their coordinates as k-dimensional trees. The distance to this axis and to its partner axis is calculated. If the distance between the focus-associated axis and its neighbour axis at this location is greater than the distance from the focus centroid to the same location in the neighbour axis, then the focus is judged to be between the axes. If this is not the case then it is outside the axes (Figure 7A-D). As with other scripts, the “Focus_position_relative_to_individually_traced_axis_pairs.py” script for batch analysis is set up to run on example data, assuming this data and necessary modules are located on the Desktop. These paths should be changed if necessary. The name of the labelmap image to be analysed should be defined as well as the positioning of the focus staining channel in the original fluorescence image, and a “Foci_name” you wish to use to identify the output data. Foci can also be filtered by a maximum and minimum area filter.

We have previously used this type of analysis to investigate focal localisations of proteins which could be involved in driving and/or maintaining interactions between axes in mutants unable to form a synaptonemal complex [19, 20]. As a further demonstration of the data generated using this method we ran the analysis on RAD51 foci segmented from the *Sycp1*^-/-^ image dataset previously analysed for axis proximity. Plotting the distribution of foci away from the axes shows that RAD51 focus positioning is a bimodal, typically falling onto one side of the axis or the other, but with a large bias towards foci being positioned between the traced chromosomes (Figure 7E). Foci are positioned a similar distance from the axis with both orientations (between axis: median 78 nm, outside of axes: median 73 nm).

It has been reported from qualitative manual analysis of SIM images in wild type mice that recombination foci formed by RPA move from being located along the centre of chromosome axes in early meiosis to moving away from the axis into the space between paired axes after chromosome synapsis [25], possibly reflecting a structural change in the recombination intermediate as recombination proceeds. RPA binds to resected ssDNA at sites of double stranded DNA breaks during early meiosis, which are also bound by RAD51. RPA additionally appears to bind D-loop DNA after strand invasion [26], an event potentially located between chromosomes and possibly reflected in the latter positioning of RPA foci between homologous axes [25]. Therefore it might be expected that the earlier localisation of RPA tightly along chromosome axes would be shared by RAD51, however this is not a pattern seen in our data, where rather than being positioned along the axis centre, RAD51 foci in asynapsed *Sycp1*^-/-^ spermatocytes are typically adjacent to the axes, most commonly being positioned between the paired axes (Figure 7E). This difference in localisation may reflect a distinction between RPA and RAD51 recombination foci or the cells analysed in the two analyses (*Sycp1*^-/-^ and WT), but the ability to objectively address this question highlights the value of our quantitative approach.

The preferential localisation of RAD51 between paired chromosome axis in *Sycp1*^-/-^ spermatocytes potentially reflects engagement of RAD51-containing single-stranded DNA ends in inter-homolog repair events. Therefore, the frequency of these inter-homolog repair events could potentially be associated with the distance between paired axes. Indeed, data generated from our tool enable us to plot axis proximity at locations where RAD51 is present either between or outside axis pairs. By plotting the ratio of RAD51 foci orientation between and outside axes, for axes with different levels of separation, we see an enrichment of foci between axes which begins when axes are 800-1000nm apart. This enrichment steadily grows until axes are 250nm apart (Figure 7F). Beyond this proximity the enrichment of RAD51 foci between paired axes steadily declines until below 100nm axis separation when foci are more likely to be positioned outside the axis pairs than between them (Figure 7F). The 250nm axis proximity where orientation of RAD51 foci between axes peaks is strikingly similar to the 193nm separation of traced axial element separation we measured in synapsed wild type spermatocytes (Figure 6B). This may therefore reflect an optimal distance for homologue engagement and procession of recombination, a distance which is maintained by the synaptonemal complex in wild-type meiosis to facilitate this process. Furthermore, if inside:outside bias in focus positioning does reflect inter-homologue intermediates, then these intermediates seem to be able to link axes separated by as much as 800-1000nm, and could posentially initiate alignment and subsequent synapsis of homologous chromosomes. Together the computational tools we share here make it possible to reveal and investigate diverse complex patterns such as these, facilitating the accumulation of a rich and detailed understanding of meiotic processes.

## Notes

Note 1. Note that when using OneDrive, a OneDrive Desktop is the system default. This location is not the default for our program. Rather, it is the user Desktop.

Note 2. We recommend saving output metadata folders in the same folder as the images being analysed (as in the Widefield_sample_data example). The macro will find existing metadata using this structure. However, because this existing data is used to populate some parameters used by the macro, to test full functionality of the script on the test image, first copy the image to a different directory and run there.

Note 3. If additional channels are to be analysed, remember to assign the output with a different name to existing focal/axial output so files are not overwritten.

Note 4. Although we routinely use this option to distinguish autosomes and sex chromosomes during analysis of mammalian meiotic chromosome spreads, this option can also be used to separate images into two compartments based on any morphological or immunostaining features in the image.

Note 5. An axes mask and focus mask must be set for the macro to work. It does not with each input independently.

Note 6. It is possible to build up masks and measurements for multiple axial and focal staining patterns within a single image by re-running the macro and changing the Axes analysis title and/or Foci analysis title as well as the associated channel number. This data will be added to any existing output metadata.

Note 7. The Gaussian-filtered image can be inspected at this point by adjusting the threshold so the mask is removed.

Note 8. If Autosome analysis has been previously selected then choose the ROI for the full axial mask including sex chromosomes. The macro will recognise the autosome analysis selection in the data and run measurements to this selected ROI mask, and the next ROI in the ROI manager assuming this is the autosomal version of this mask.

Note 9. If “Autosome analysis” was selected earlier then the removed sex chromosomes will reappear here. This provides an opportunity to also optimise the sex chromosome skeletons so measurements relating to them can be calculated by subtracting the autosomal from the total measurements.

Note 10. If both members of a pair of axes are being traced, name these traces with the same number and differing letters e.g. 1a and 1b. This nomenclature is recognised in other tools herein for quantifying inter-axis distances and orientated focus positioning relative to axis pairs.

Note 11. For tracing resolved super-resolution axes it can be beneficial to reduce the “Enable Snapping within XY” value in SNT to allow separation of proximal axes. Alternatively, to generate a single trace for a pair of axes the image can be processed using a Gaussian filter to blur the axes together.

Note 12. During this analysis the inter-focus measurements generated are two-dimensional, these are not one-dimensional measurements of proximity along the chromosome axis.

Note 13. It may be necessary to adjust the parameters of “Enable Snapping within XY” within SNT. Applying the “Sharpen” function to the axial image can also benefit tracing.

Note 14. The closest pixel in the partner axis may not always correspond to the same position along the chromosome.

---

Supplementary Figure 1. RAD51 focus positioning on asynapsed pachytene-like autosomes in *Sycp1*^-/-^ spermatocytes.

Barplot displaying the percentage of all RAD51 foci measured which are positioned in each 10% of axis length from the centromere. Images and axes included in this analysis are identical to those in Figure 6.

## Acknowledgements

We wish to thank all the members of the Institute of Genetics and Cancer Advanced Imaging Resource team, for all their microscopy-related assistance, in particular Laura Murphy for sharing image-analysis advice. We are also grateful to Willy Baarends (Erasmus MC, Rotterdam) for sharing Sycp1^-/-^ spermatocyte spreads, and C. James Ingles (University of Toronto, Canada) for sharing RPA antibody.

