## Supplementary figures and images for "Computational Tools for the Analysis of Meiotic Prophase I Images"

### Supplemental Figure 1

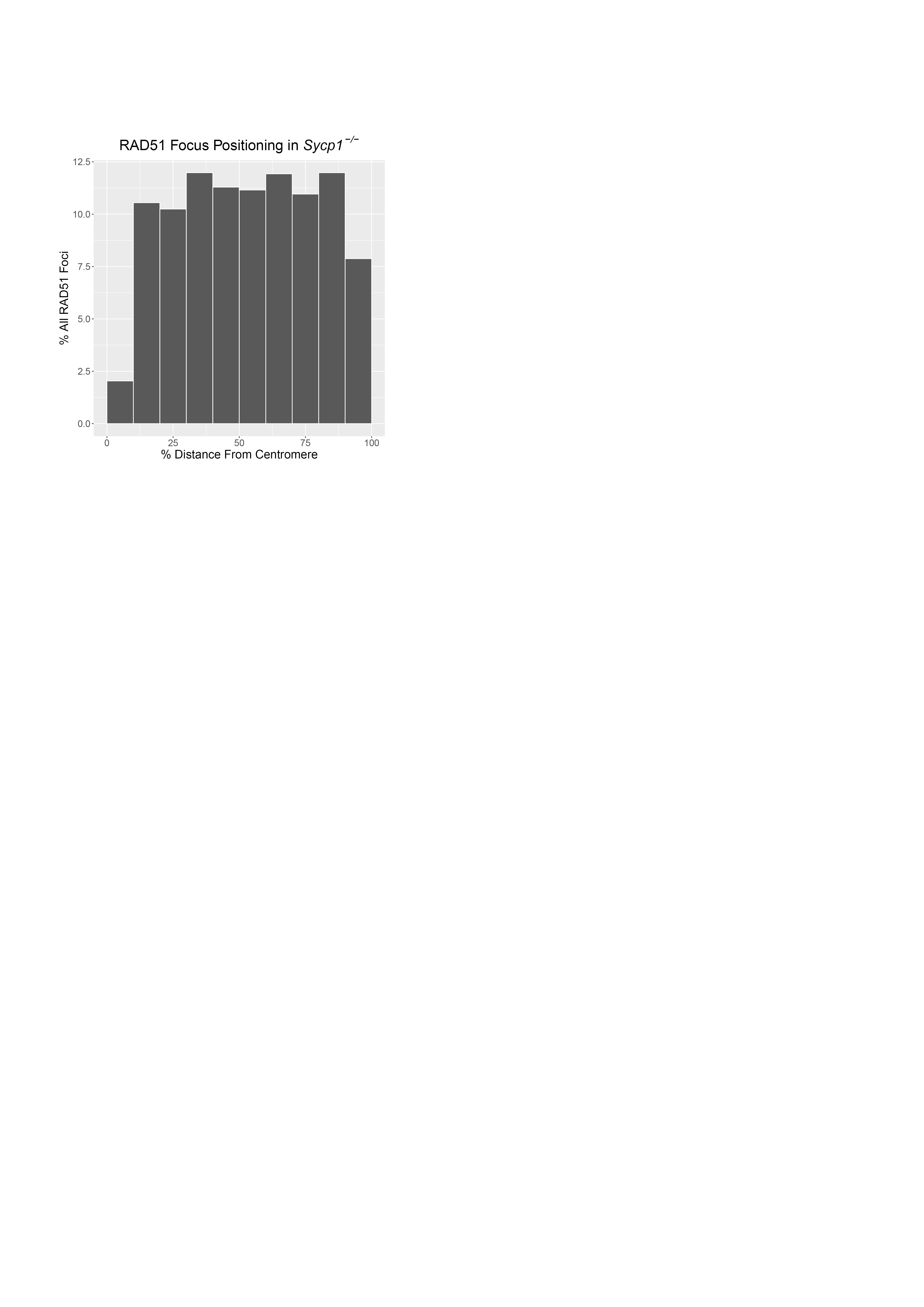
